# Modulatory upregulation of an insulin peptide gene by different pathogens in *C. elegans*

**DOI:** 10.1101/238568

**Authors:** Song-Hua Lee, Shizue Omi, Nishant Thakur, Clara Taffoni, Jérôme Belougne, Ilka Engelmann, Jonathan J. Ewbank, Nathalie Pujol

**Affiliations:** CIML, Centre d’Immunologie de Marseille-Luminy, Aix Marseille Université UM2, INSERM U1104, CNRS UMR7280, 13288 Marseille, France

## Abstract

When an animal is infected, its innate immune response needs to be tightly regulated across tissues and coordinated with other aspects of organismal physiology. Previous studies with *Caenorhabditis elegans* have demonstrated that insulin-like peptide genes are differentially expressed in response to different pathogens. They represent prime candidates for conveying signals between tissues upon infection. Here, we focused on one such gene, *ins-11* and its potential role in mediating cross-tissue regulation of innate immune genes. While diverse bacterial intestinal infections can trigger the up-regulation of *ins-11* in the intestine, we show that epidermal infection with the fungus *Drechmeria coniospora* triggers an upregulation of *ins-11* in the epidermis. Using the *Shigella* virulence factor OpsF, a MAP kinase inhibitor, we found that in both cases, *ins-11* expression is controlled cell autonomously by p38 MAPK, but via distinct transcription factors, STA-2/STAT in the epidermis and HLH-30/TFEB in the intestine. We established that *ins-11*, and the insulin signaling pathway more generally, are not involved in the regulation of antimicrobial peptide gene expression in the epidermis. The up-regulation of *ins-11* in the epidermis does, however, affect intestinal gene expression in a complex manner, and has a deleterious effect on longevity. These results support a model in which insulin signaling, via *ins-11*, contributes to the coordination of the organismal response to infection, influencing the allocation of resources in an infected animal.

## Introduction

Infection of *Caenorhabditis elegans* induces broad changes in gene expression (reviewed in ^1, 2^). These are characterized by pathogen-specific responses and more generic changes ^3^. We previously noted among the genes induced by infection with the fungi *Drechmeria coniospora* an overrepresentation of genes repressed after infection by the bacterial pathogens *Serratia marcescens, Enterococcus faecalis* or *Photorhabdus luminescens*. Conversely, there was an overrepresentation of genes up-regulated upon infection by each of these bacterial pathogens among the genes down-regulated upon infection with *D. coniospora* ^4^ Since *D. coniospora* infects via the epidermis, while the bacterial pathogens colonize the nematode intestine, one possible interpretation of these results is that there is a communication between tissues. Thus, when immune defenses are activated in the epidermis, they would be repressed in the intestine and vice versa. In a parsimonious model, fungal infection of the epidermis or bacterial infection of the intestine would provoke an increase in the expression of a signaling molecule that would then modulate gene expression in other tissues.

There are countless instances of such non-cell autonomous control of gene expression across all multicellular species. A number have been characterized in the context of the response of *C. elegans* to biotic and abiotic insults (reviewed in ^5^). To give one recent example, activation of a conserved p38 MAPK pathway in the intestine by rotenone is associated with protection from neurodegeneration ^6^, conceivably because induction of the mitochondrial unfolded protein response in the intestine represses the expression of the antimicrobial peptide gene *nlp-29* ^7^, which regulates dendrite degeneration in aging and infection ^8^. But, as is commonly the case, the molecular nature of the signal mediating this cross-tissue control has not yet been established. One prominent exception to this is the complex network of insulin signaling. With 40 genes encoding insulin-like peptides ^9^, acting through a common conserved pathway to regulate the activity of the FOXO transcription factor DAF-16 and thereby the expression of hundreds of target genes across all tissues ^10^, insulin signaling influences many if not all aspects of *C. elegans* physiology, including diapause, longevity, stress resistance and fertility ^9^. It has been demonstrated to play a role in innate immunity too. Specifically, the insulin-like peptide INS-7 from neurons has been shown to inhibit DAF-16 signaling in the intestine and thereby negatively-regulate pathogen resistance ^11^. Interestingly, infection with *Pseudomonas aeruginosa* provokes an increase in neuronal *ins-7* expression, leading to the down-regulation of putative immune defense genes in the intestine ^12^. And recently, a role in learning aversive behavior has been proposed for insulin-like peptide INS-11 upon infection by *P. aeruginosa* ^13^. In this study, we investigated the induction of *ins-11* caused by different infections and propose a non-cell autonomous role for *ins-11* in cross-tissue communication.

## Results and Discussion

### Expression of insulin peptides upon infection

Given their known roles in cross-tissue communication and in innate immune regulation, insulin-like peptides are prime candidates for mediating the reciprocal control of epidermal and intestinal gene expression seen following infection of *C. elegans*. We therefore first reviewed data from our previous studies to identify insulin-like genes that are induced both by fungal and bacterial pathogens. Infection of *C. elegans* by *D. coniospora* provokes a marked increase in the expression of 4 insulin-like genes, *ins-7, ins-11, ins-23* and *ins-36* ^4^ Among them, although *ins-7* expression is induced by *P. aeruginosa* ^12^, it did not figure among the list of genes differentially regulated upon infection by the bacterial pathogens *S. marcescens*, *E. faecalis* or *P. luminescens*. And while the expression of *ins-23* and *ins-36* was decreased upon infection with *S. marcescens*, for *ins-11* it was increased, as it was also following infection with *E. faecalis* and *P. luminescens* ^4^. Further, other investigators have reported an induction of *ins-11* expression after infection with *Bacillus thuringiensis* ^14^, *P. aeruginosa* ^13, 15, 16^, *Staphylococcus aureus* ^17^ and the microsporidian species *Nematocida parisii* ^18^. We therefore focused our attention on *ins-11* and its potential role in mediating a cross-tissue regulation of innate immune genes.

### *hlh-30* dependent upregulation of *ins-11* in the intestine upon bacterial infection

As a first step in the characterization of *ins-11*, we verified its expression upon infection by *S. marcescens*. By qRT-PCR we observed a clear induction of *ins-11* gene expression, as we did for the C-type lectin gene *clec-67*, a putative defense gene that is expressed in the intestine and is up-regulated by infection by diverse bacteria ^3^ (Fig. 1A). It has previously been shown that following *S. aureus* or *P. aeruginosa* infection, *ins-11* expression requires the basic helix-loop-helix transcription factor HLH-30 ^13, 17^ In the case of infection by *S. marcescens, ins-11* expression was also dependent upon *hlh-30* (Fig. 1A). As previously reported ^19, 20^, a strain carrying an *ins-11* transcriptional reporter construct (*ins-11p::gfp*) exhibited only a low constitutive fluorescence in adults, in the nerve ring, labial neurons and most obviously in the intestine (Fig. 1B). When this strain was infected by *S. marcescens*, a clear increase in GFP expression was observed in the intestine (Fig. 1C). Interestingly, the same type of induction in the intestine was also recently reported upon infection by *P. aeruginosa* ^13^. This suggests that diverse bacterial intestinal infections can trigger the up-regulation of *ins-11* in the intestine.

**Figure 1.**
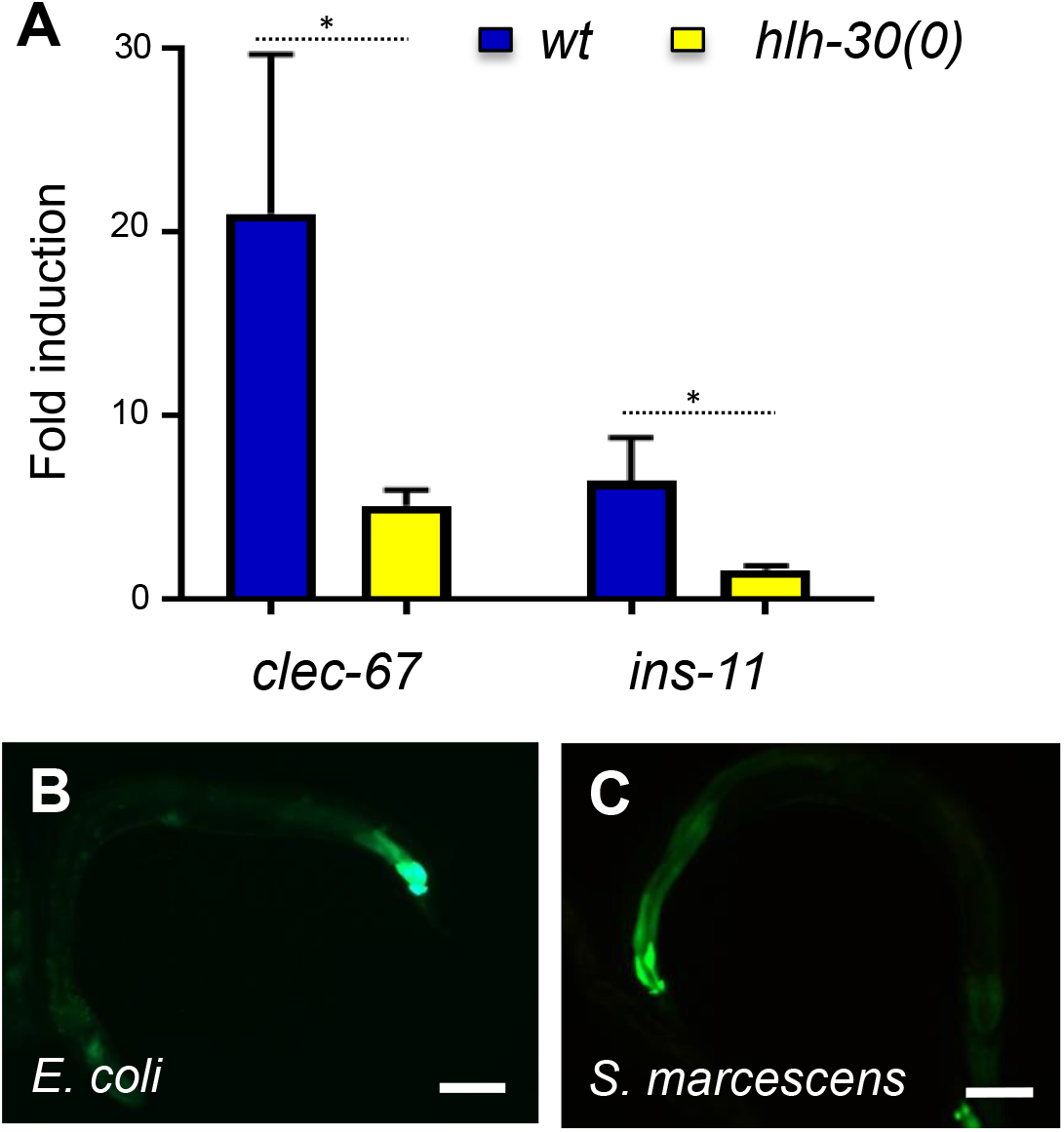
*S. marcescens-induced ins-11* expression in the intestine requires *hlh-30*. (**A**) Relative induction of gene expression as measured by qRT-PCR at 24 hours post infection with *S. marcescens* in wild-type worms (blue) or in *hlh-30* mutants (yellow), * p value < 0.05. (**B, C**) Representative fluorescence images showing a strain carrying the *ins-11p::GFP* transcriptional reporter under the normal conditions, *E. coli* (**B**) or infected with *S. marcescens* for 24 h (**C**). Scale bar 50 μm. For a quantification of GFP levels, see below (Fig. 3C).

### *sta-2* dependent upregulation of *ins-11* in the epidermis upon fungal infection

We then assayed the expression of *ins-11* upon infection by *D. coniospora* by qRT-PCR, and confirmed its strong induction, of the same order of magnitude as that of the well-characterized antimicrobial peptide gene *nlp-29* (Fig. 2A). Infecting a strain carrying an *nlp-29p::GFP* reporter with *D. coniospora* provokes GFP expression in the epidermis, particularly at the perivulval cells when spores adhered to this region of the adult cuticle ^21^. A very similar pattern was seen when we infected worms carry the *ins-11p::GFP* reporter (Fig. 2B, C). It is notable that epidermal expression of *ins-11* has not previously been reported ^19^. Strong GFP expression was also observed in the epidermis when the *ins-11p::GFP* reporter was transferred into a strain that expresses a constitutively active Gα protein, GPA-12*, in the adult epidermis (Fig. 2D); *nlp-29p::GFP* expression is similarly elevated in the GPA-12* background ^22^. GPA-12 acts upstream of the PMK-1 p38 MAPK cascade to regulate *nlp-29* ^23^. Our result suggests that *ins-11* expression in the epidermis might be controlled in the same manner as *nlp-29*. While *hlh-30* is dispensable for *nlp-29* expression following *D. coniospora* infection (^7^ and results not shown), the STAT-like transcription factor gene *sta-2* has been shown to have a specific function in the regulation of antifungal defenses in the epidermis, downstream of the GPA-12/PMK-1 pathway ^24^ Similarly, we found the induction of *ins-11* upon infection by *D. coniospora* to depend very largely upon *sta-2* when we assayed the level of the endogenous gene by qRT-PCR (Fig. 2A) or the fluorescence level of the reporter strain (Fig. 2B). Our results therefore suggest that *ins-11* expression is triggered in infected tissues, and regulated in the epidermis and intestine by distinct transcription factors.

**Figure 2.**
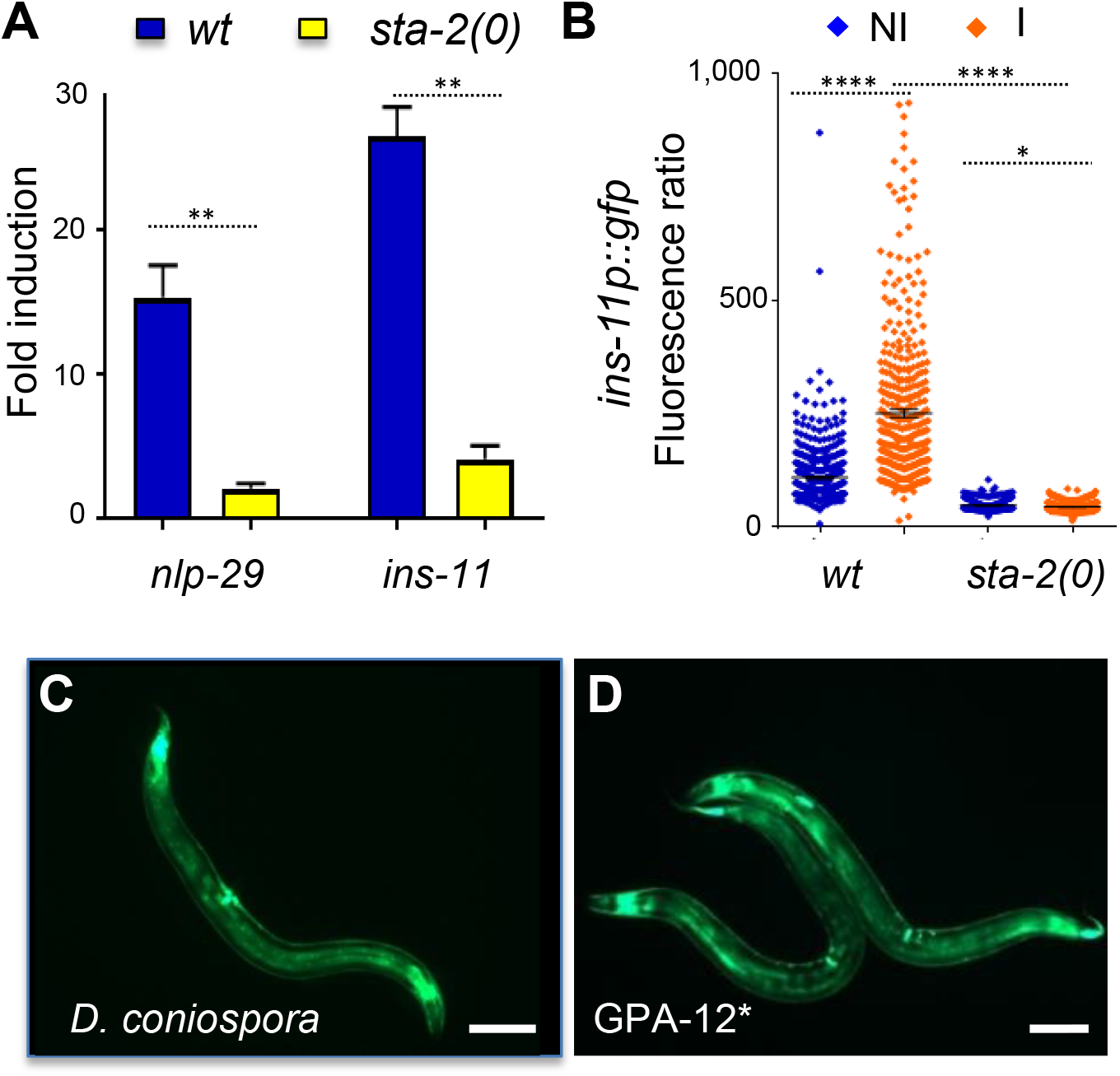
*D. coniospora-induced ins-11* expression in the epidermis requires *sta-2*. (**A**) Relative induction of gene expression as measured by qRT-PCR at 8 hours post infection with *D. coniospora* in wild-type worms (blue) or in *sta-2* mutants (yellow). ** p< 0.01 (**B**) Quantification of relative green fluorescence in a strain carrying the *ins-11p::GFP* transcriptional reporter in non-infected (NI) or after 16 h of infection (I) with *D. coniospora* (blue and orange symbols, respectively) in a wild-type or *sta-2* mutant background (n = 496, 399, 381, 532), **** p value < 0.0001. (**C, D**) Representative fluorescence images showing a strain carrying the *ins-11p::GFP* transcriptional reporter infected with *D. coniospora* (**C**) or in a strain expressing the constitutively active Gα protein, GPA-12* (**D**). Scale bar 50 μm.

### Cell autonomous inactivation of p38 MAPK PMK-1

To investigate the regulation of *ins-11* in different tissues, we made use of the *Shigella flexneri* protein effector OspF that irreversibly dephosphorylates p38 ^25^ since this offers a straightforward way to affect signaling in a temporally-controlled and tissue-specific manner. OspF is a dual-specificity phosphatase and also dephosphorylates ERK MAPKs. While the nematode ERK pathway involving the MAPK MPK-1 has been shown to be important for the rectal epithelial cell swelling response to *Microbacterium nematophilum* ^26^ and for the swelling response to *Staphylococcus aureus*, it is dispensable for the intestinal transcriptional response ^27^. The ERK pathway is not involved in the response to *D. coniospora* either ^7, 28^, so if OspF has an effect on infection-induced gene expression, it is expected to be solely via the p38 MAPK pathway. Since we have previously observed unexpected changes in gene expression patterns when we interfered with p38 signaling in a single tissue ^7^, we used the transcriptional reporters described above to assay the effect of tissue-specific inhibition of MAPK activity. Expression of OspF in the intestine, under the control of the *vit-6* promoter, had little or no effect on the induction of *nlp-29p::GFP* expression that normally accompanies infection by *D. coniospora*. When OspF was expressed under the control of an adult epidermis-specific promoter (*col-19p*), on the other hand, there was a very marked reduction in *nlp-29p::GFP* expression (Fig. 3A, B). These results are consistent with our previous findings that *nlp-29* is controlled in the epidermis in a cell autonomous manner by the p38 MAPK PMK-1 ^29^. Expressing OspF in the intestine did abrogate *ins-11p::GFP* expression in the intestine following infection with *S. marcescens* expression, while its expression in the epidermis strongly inhibited *ins-11* reporter gene expression upon *D. coniospora* infection (Fig. 3C, D). Together, these results suggest that *ins-11* expression is controlled cell autonomously by p38 MAPK signaling both in the epidermis and in the intestine, via distinct downstream transcription factors (STA-2 and HLH-30).

**Figure 3.**
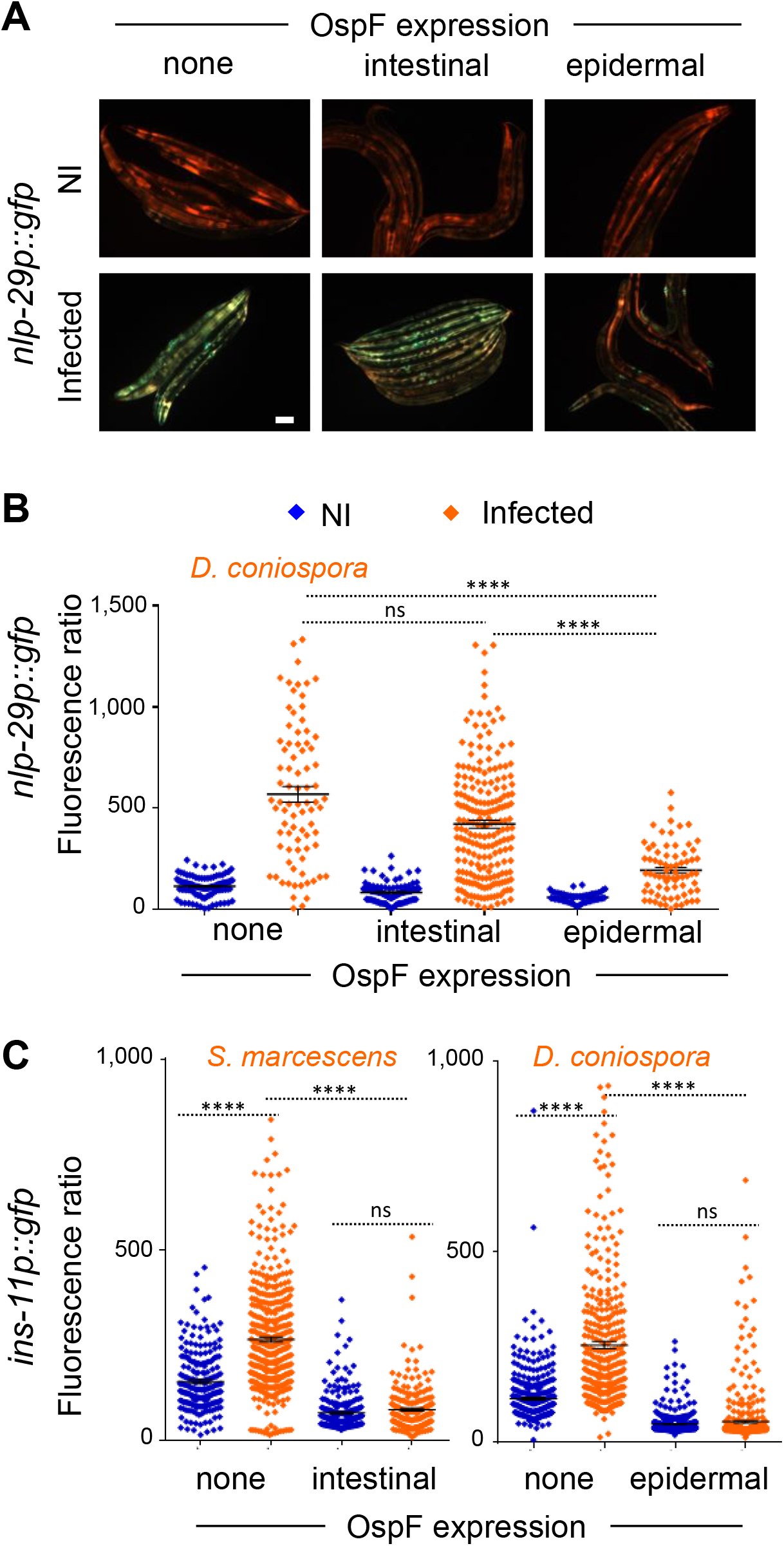
Infection-induced *ins-11* expression requires p38 MAPK signaling. (**A**) Representative fluorescence images of a strain carrying the *nlp-29p::GFP* transcriptional reporter alone (none, left hand panels), or in combination with *frSi14[vit-6p::OspF]* (intestinal, middle panels) or *frSi15[col-19p::OspF]* (epidermal, right panels), in non-infected (NI) worms or after 16 h infection with *D. coniospora* (Infected; upper and lower panels, respectively). Green and red fluorescence are viewed simultaneously. Scale bar 50 μm. (**B**) Quantification of relative green fluorescence in the same strains, in non-infected (NI, blue) worms or after 16 h of infection with *D. coniospora* (I, orange), n = 105, 81, 140, 212, 59, 73. (**C**) Quantification of relative green fluorescence in a strain carrying the *ins-11p::GFP* transcriptional reporter on its own (none), together with *frSi14* (intestinal), or *frSi15* (epidermal) in control populations (blue symbols) or after infection (orange symbols) with *S. marcescens* (left panel, n = 222, 444, 167, 181) or *D. coniospora* (right panel, n = 496, 399, 584, 509). **** p value < 0.0001 and ns p > 0.05.

### No cell autonomous role of *ins-11* or *daf-16* upon infection

Zou et al. reported that infection by *D. coniospora* leads to nuclear translocation of the FOXO transcription factor DAF-16 ^30^. With the *D. coniospora* strain ^31^ used in this and all our previous studies (ATCC 96282), and the DAF-16::GFP reporter protein used by Zou et al., we failed, however, to observe any such translocation. DAF-16 also remained cytoplasmic in the strain expressing GPA-12* in the epidermis that mimics *D. coniospora* infection (see below; results not shown). Further, fungal infection did not interfere with the normal cytoplasm-nucleus shuttling of DAF-16 associated with transient heat-shock ^32^ (Fig. 4A). Thus contrary to the results of Zou *et al*. ^30^, we have no evidence to suggest that infection by *D. coniospora* leads to nuclear translocation of DAF-16. As we have not been able to procure the strain used by Zou *et al.*, nor garner any information about it, the reason for the discrepancy is unclear.

**Figure 4.**
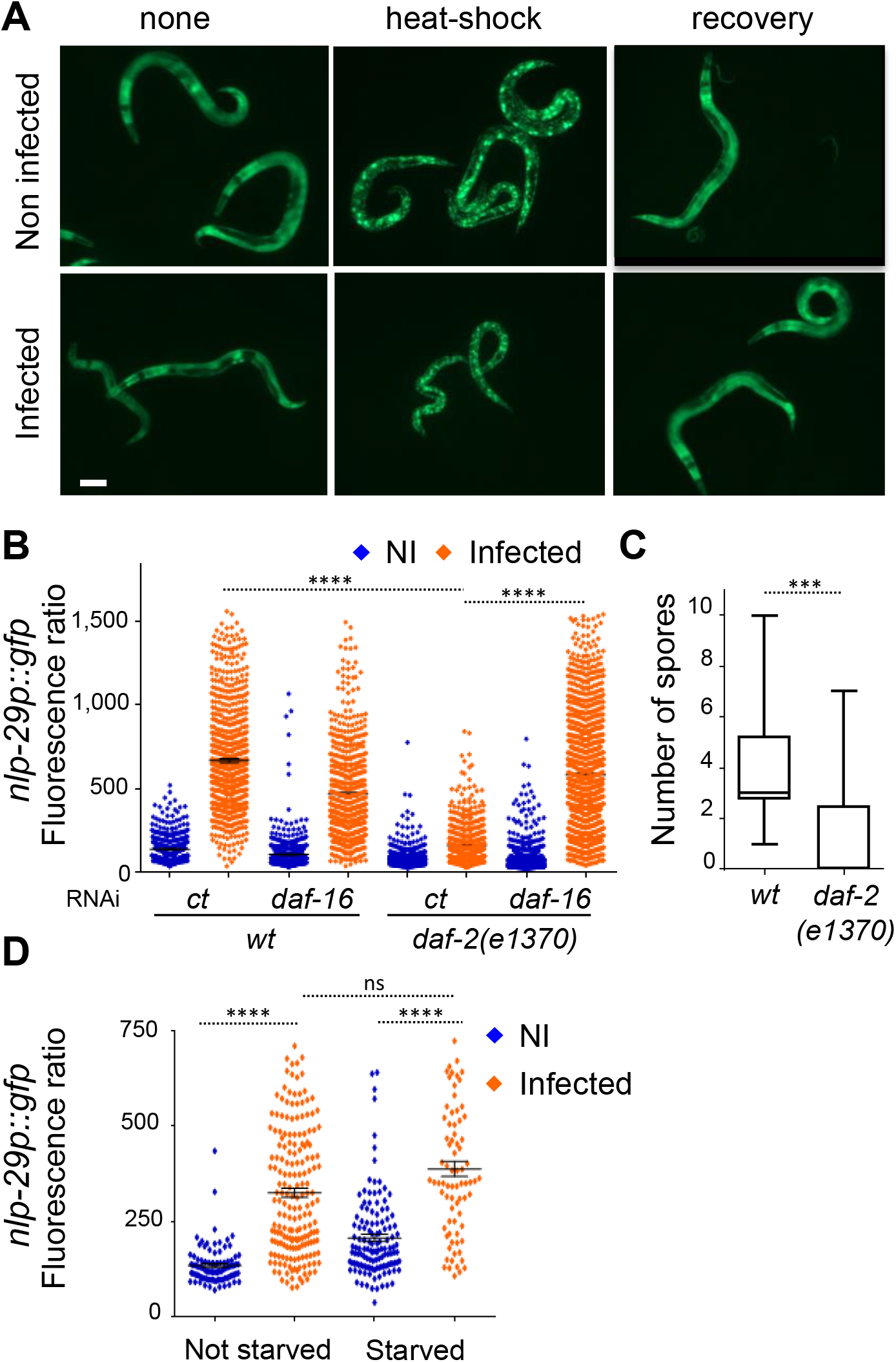
Epidermal *nlp-29* expression is independent of insulin signaling. (**A**) Representative fluorescence images of TJ356 worms expressing a DAF-16::GFP fusion protein under normal culture conditions at 25 °C (none, left hand panels), after 30 minutes at 37 °C (heat-shock, middle panels) or after a subsequent hour at 25 °C (recovery, right panels), without infection (Non-infected, upper panels) or after 13 h of infection with *D. coniospora* (Infected, lower panels). Scale bar 50 μm. (**B**) Quantification of relative green fluorescence in the wild type and the *daf-2(e1370)* mutant strain subjected to control (ct) or *daf-16* RNAi, in non-infected (NI, blue) worms or after 16 h of infection with *D. coniospora* (I, orange), n = 500, 728, 621, 595, 666, 604, 1151, 1210. (C) Quantification of the number of spores attached to wild type and *daf-2(e1370)* young adult worms, n =18 and 13, respectively. (**D**) Quantification of relative fluorescence in wild type worms carrying the *nlp-29p::GFP* transcriptional reporter in non-infected worms or after infection for 6 h with *D. coniospora* (blue and orange symbols) after normal ad libitum feeding, or 14 h in the absence of food (starved; n = 98, 191, 129, 80). **** p value < 0.0001, *** p < 0.001, ns p > 0.05.

We further tested *daf-2(ts)* mutant worms in which DAF-16 is constitutively active at the restrictive temperature (25 °C) with a predominantly nuclear localization^32, 33^. In the *daf-2* mutant background, there was a marked reduction of the induction of the antimicrobial peptide reporter gene *nlp-29p::GFP* compared to wild-type worms that was suppressed by *daf-16* inactivation (Fig. 4B). *daf-2* mutants have an altered cuticle ^34^. Changes in the surface coat or cuticle can affect the binding of fungal spores ^7, 35^. We found that the reduced induction of *nlp-29p::GFP* correlated with decreased binding of *D. coniospora* spores to the *daf-2* mutant cuticle (Fig. 4C). These results serve as a caveat for studies using these mutants. To circumvent the changes in cuticle in *daf-2* mutants, we used starvation to drive DAF-16 into the nucleus ^32^ in young adult worms. In this case, the induction of *nlp-29p::GFP* expression upon infection was normal (Fig. 4D). Taken together, our results suggest that the insulin signaling pathway is unlikely to impinge directly on the cell autonomous intracellular p38 MAPK pathway known to control *nlp* antimicrobial peptide gene expression.

### Activation of GPA-12 in the epidermis mimics *D. coniospora* infection at the transcriptional level

To determine whether *ins-11* might play a role in innate immunity by regulating the expression of defense genes other than *nlp-29*, we conducted an exploratory transcriptomic study using RNA-seq to compare the transcriptional profiles of wild type, and GPA-12*-expressing animals, with and without *ins-11*. Expression was detected for 19,845 genes in at least one condition. Principle component analyses neatly separated the two “treatments”, loss of *ins-11* function and gain of function of *gpa-12*, with the first 2 components accounting for most of the variability (49 *%* and 27 %; Table S1). As expected, class enrichment analysis (^7^; bioinformatics.lif.univ-mrs.fr/YAAT; Thakur et al. in preparation) showed a very substantial overlap (117/252 genes; 46 %; p < 10^−67^) between the list of genes up-regulated in GPA-12* worms (relative to N2) and the list of genes up-regulated upon *D. coniospora* infection ^4^, including *ins-11* (> 25-fold), and more than 20 genes that potentially encode AMPs ^36^ (with 8 *cnc*, 5 *fip* and *fipr*, 7 *nlp* genes; Table S1). It also identified a number of other categories previously noted to be enriched for innate immune genes. This includes genes regulated by *nhr-25* and *acs-3* ^37^, *sam-10* ^38^ or *dapk-1* ^39^, induced by infection with *Burkholderia pseudomallei* ^40^ and in a series of mutants with an alteration of their cuticle and/or osmotic balance (*osm-7, osm-8, osm-11*) ^41^. As expected, there was also a very significant enrichment (p < 10^−28^) for genes characterized as being expressed in the epidermis ^42^ (Table S1). Interestingly, regarding the genes down-regulated both by infection and in the GPA-12* background, the overlap was proportionally smaller (104/653) although still significant (p < 4.5 × 10^−4^), and there were comparatively fewer epidermally-expressed genes (33/104 vs 61/117). One could speculate that *D. coniospora* might be able to down-regulate host gene expression through the action of its many candidate secreted virulence factors ^43^. These analyses nevertheless validate the use of strains carrying *col-19p::GPA-12\** as a model for the inductive part of the epidermal innate immune response.

### A potential non cell-autonomous role of *ins-11* in transcriptional regulation

Since the expression of *ins-11* is modulated by the GPA-12/p38 MAPK cascade that is central to the transcriptional response to *D. coniospora* infection, any gene that is a target of *ins-11* regulation should also appear to be regulated in a concordant manner by *D. coniospora* infection and the GPA-12/p38 MAPK cascade. Comparing the list of genes that were up regulated by expression of GPA-12* in the wild-type versus *ins-11(0)* background, we observed 130 genes in common (34%) that are thus targets of the p38 cascade, regulated in a manner independent of *ins-11* (Fig. 5A, Table S2). Consistent with our results with the *nlp-29p::GFP* reporter gene above, these included *nlp* and *cnc* genes, which encode secreted AMPs, and other genes known to be expressed in the epidermis (as mentioned above, ^44^). In line with this, among the 8 Gene Ontology (GO) terms returned by class enrichment analysis (^7^; Thakur et al. in preparation), the one with the highest probability was “defense response to fungus” (GO:0050832; Fig. 5A; Table S2). On the other hand, 122 genes were found to be up-regulated only in the wild-type (32%) and their expression is thus potentially dependent on *ins-11*. Interestingly, the list included genes known to be up-regulated in the intestine by infection (like *irg-4* ^45^, *irg-5* ^46^ and *kreg-1* ^47^ or abiotic stress like glycosyltransferase (*ugt*) or cytochrome P450 (*cyp*) genes; the associated GO terms included “innate immune response” (GO:0045087). A last group of 127 genes (34%) were only found to be up-regulated in the *ins-11(0)* mutant background, so these are potentially repressed by *ins-11* upon GPA-12 activation. The only significantly enriched GO term, “defense response to Gram-positive bacterium”, (GO:0050830) reflected the many intestinal genes differentially regulated by *Bacillus thuringiensis* (such as *lys-7;* see below), but the list also included genes expressed in the epidermis such as collagen, cuticulin, astacin and patched-like genes ^42, 44^

**Figure 5.**
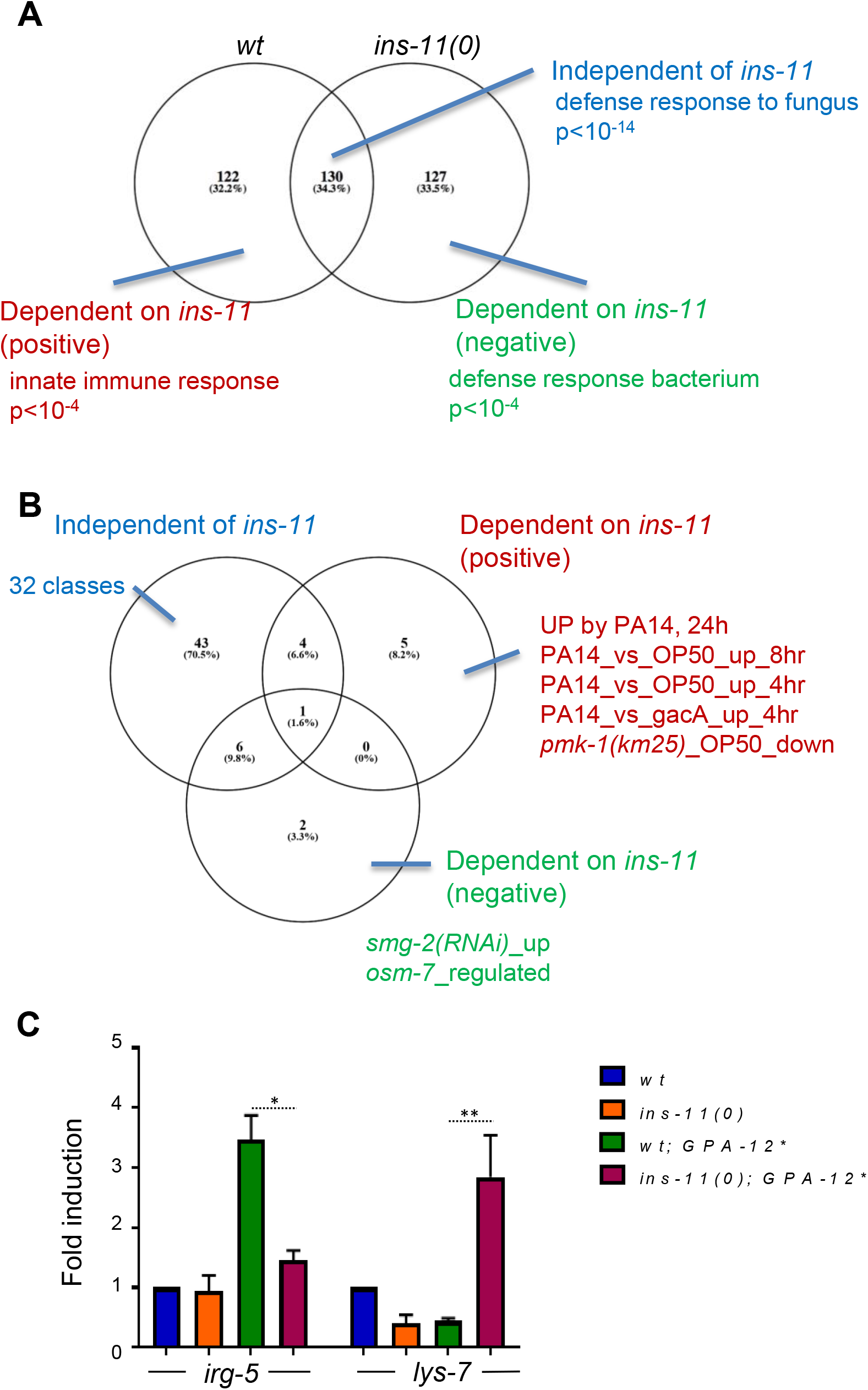
(**A**) The genes up-regulated by expression of GPA-12* in the wild type or the *ins-11(0)* background are analyzed in a Venn diagram. The overlap includes genes that are up-regulated by GPA-12* in an *ins-11* independent manner. The top-ranking enriched GO terms associated with each group are shown. (**B**) The 3 gene lists defined above were analyzed for enrichment of gene classes (p < 10^−10^). The classes are indicated when they are not too numerous; the full list is provided in Table S2. (**C**) Relative expression of *irg-5* and *lys-7* in wild-type and *ins-11* backgrounds, with and without the expression of GPA-12*. ** p value < 0.01, * p < 0.05.

Beyond the current GO terms for *C. elegans* genes, the analysis identified enrichment in the common gene set (*ins-11*-independent) for classes associated with epidermal and fungal innate immune defense, while intestinal bacterial immune classes were specifically associated with the ins-dependent gene set. Prominent among the significantly enriched classes specific to the genes positively regulated by *ins-11* were several *P. aeruginosa* up-regulated genes (p < 10^−10^ Fig. 5B, Table S2). We picked two genes known to be involved in intestinal defense, *irg-5* ^46, 48^ and *lys-7*^12, 49–51^. By qRT-PCR, we confirmed that their expression was dependent on *ins-11*, in line with the RNAseq data, being positively and negatively regulated by *ins-11*, respectively (Fig. 5C). These data suggest that the up-regulation of *ins-11* in the epidermis can affect innate immune gene expression in the intestine, in a complex manner.

### *ins-11* expression in the young adult epidermis affects longevity

As mentioned above, *ins-11* is not strongly expressed under normal laboratory conditions and only mild developmental and physiological phenotypes have been observed in the *ins-11* mutant: increased fat accumulation in the adult ^52^ and an increase in the number of eggs laid ^13^. On the other hand, *ins-11* is expressed in the epidermis upon fungal infection, and under these conditions its loss might be expected to have more marked consequences. We therefore decided to assay the effect of the loss of the *ins-11* in a strain carrying the gain of function Gα protein (GPA-12*) mentioned above that mimics infection. Under these conditions, we observed a small but significant increase in the rate of egg laying upon loss of *ins-11*. Since this is similar to the increase recently reported in the wild-type background ^13^, it indicates that *ins-11* expression in the epidermis does not have a major influence on this phenotype (Fig. 6A). We also noted a small but significant difference in the size of adult GPA-12* worms upon loss of *ins-11*, but again, as this effect was seen when comparing *ins-11* mutants and wild type worms, this phenotype is a not primarily a consequence of expression of *ins-11* in the adult epidermis (Fig. 6B & Fig. S1).

**Figure 6.**
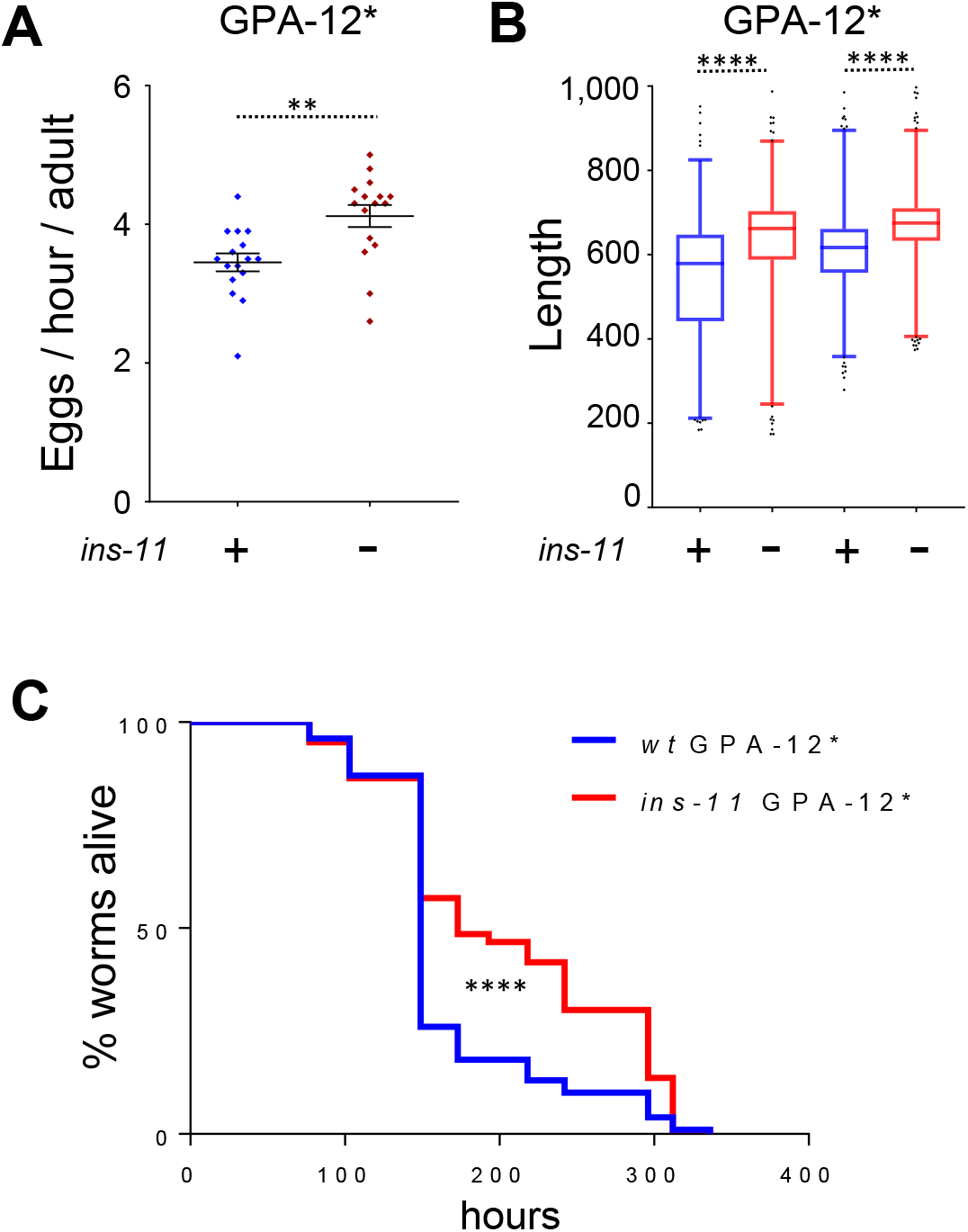
*ins-11* affects egg laying, adult size and longevity. The effect of *ins-11* was determined under conditions when its expression is elevated in the adult epidermis by comparing worms expressing GPA-12* in the wild-type (blue) or *ins-11* backgrounds (red) backgrounds. (**A**) The number of eggs laid par adult per hour is presented as an average for 15 independent plates (total number of F1 worms analyzed are 3,177 and 3,836, respectively). (**B**) Relative size of young adult worms from eggs laid in 2 consecutive windows of 2 hours (n = 794, 871, 876, 1025). Box and whisker plot of 1-99 percentile, **** p value < 0.0001. (**C**) Survival of young adult worms at 25 °C (n = 100 and 106, respectively), **** p value < 0.0001. The results are representative of 3 independent trials. Of note, the lifespan of the GPA-12* transgenic worms characterized here is generally shorter than the lifespan normally observed for wild-type worms in the literature (medium survival around 310 hours for example in ^52^).

Given the opposing effect of *ins-11* that we observed for immune defense gene expression (Fig. 6C, Table S2), it is difficult to predict what effect *ins-11* would have on the capacity of worms to resist infection. Indeed, no difference was reported during *P. aeruginosa* infection between the wild type and the *ins-11* mutant ^13^, and preliminary tests indicated no significant difference in the survival of adult GPA-12* worms upon loss of *ins-11* following infection with *S. marcescens* or *P. aeruginosa* (data not shown). In both cases, however, the progression of the infection is relatively rapid. In order to assay the effect of *ins-11* in a less acute context, we therefore measured the lifespan of the 2 strains in the absence of infection. We observed an increase in the longevity of the ins-11;GPA-12* worms relative to the GPA-12* controls (Fig. 6C, p value < 0.0001, median survival 149 and 173 hours for wt;GPA-12* and *ins-11*; GPA-12* strains, respectively). As no difference was observed between wild type and *ins-11* mutant worms cultured on the regular non-pathogenic OP50 bacteria ^52^, our data suggest that expression of *ins-11* in the adult epidermis has a negative effect on longevity. A previous report suggested that *ins-11* expression has a deleterious effect on the longevity of adult hermaphrodite worms in the presence of males ^53^. One can speculate that mating provokes epidermal wounding in hermaphrodites and therefore an induction of *ins-11* in the epidermis, as we observed in the GPA-12* transgenic worms. It would be interesting to test this hypothesis in the future.

Taken together, our results suggest that activation of the p38 MAPK pathway in the epidermis of *C. elegans* leads to an increase in cell autonomous defenses (e.g. AMP gene expression), but also impacts on intestinal gene expression, in part through the production of the insulin-like peptide INS-11. Although the effect of increased INS-11 levels is complex with regard the expression of individual intestinal defense genes, one consequence is that worms have a reduced longevity. We speculate that this reflects an altered allocation of energy, with resources being directed to the tissue most at threat from a pathogen. Interestingly, upon bacterial infection *ins-11* is induced in the intestine to counter balance a protective aversive behavior probably to limit the time spend out of food ^13^. Notably, in *Drosophila*, induction of an insulin-like peptide is used during development to coordinate organ growth ^54^. It would be interesting in the future to investigate whether *ins-11* serves as a sentinel during development, alerting the organism when tissue homeostasis is perturbed, by infection or other insults, thus promoting optimal growth even under conditions of stress.

## Materials and Methods

### Fungal, bacterial and nematode strains

The fungal strain used was *D. coniospora* ATCC 96282 ^43^. Bacterial strains used were *S. marcescens* strain Db10 ^55^ and *E. coli* strain OP50. All the following *C. elegans* strains were maintained on nematode growth medium (NGM) and fed with *E. coli* OP50, as described ^56^: the wild type N2, HT1714 *unc-119(ed3) III; wwEx70[ins-11p::GFP + unc-119(+)]* ^20^, FX1053 *ins-11(tm1053) II* ^52^, TJ356 *wt; zIs356[DAF-16::gfp, pRF4(rol-6(su1006))]*, JIN1375 *hlh-30(tm1978) IV* ^17^, IG1241 *sta-2(ok1860)* ^24^, IG274 wt; *frIs7[nlp-29p::gfp, col-12p::DsRed] IV* ^29^, IG571 *daf-2(e1368) III; frIs7[nlp-29p::gfp, col-12p::DsRed]* IV, IG356 *daf-2(e1370) III; frIs7[nlp-29p::gfp, col-12p::DsRed] IV*. The transgenic strains carrying *frSi2, frSi14* and *frSi15* containing respectively a single copy of *col-19p::GPA-12*, vit-6p::OspF::unc-54 3’UTR* and *col-19p::OspF::unc-54 3’UTR* were generated in this study (see Table S3 for a list of all strains).

### Constructs and transgenic lines

All the transgenic strains carrying *frIs7, frSi2* or *wwEx70* were obtained by conventional crosses and all genotypes were confirmed by PCR or sequencing. All constructs were made using Gibson Assembly (NEB Inc., MA).

*frSi2* is a single copy insertion on chromosome II at the location of the Nemagenetag *Mos1* insertion ^57^ *ttTi5605* of pNP138 (*col-19p::GPA-12\**). pNP138 was obtained by insertion of the *col-19* promoter fused to the DNA encoding an activated form of GPA-12* (with the Q205L mutation) ^22^ into the MosSCI vector pCFJ151 ^58^. It was injected (at the Lyon platform UMS3421 CNRS) into the EG6699 strain at 30 ng/μl together with pGH8 (*rab-3p::mCherry*) at 10 ng/μl, pCFJ90 (*myo-2p::mCherry*) at 2.5 ng/μl, pCFJ104 (*myo-3p::mCherry*) at 5 ng/μl, pMA122 (*hsp16.41p::PEEL-1*) at 10 ng/μl, pCFJ601 (*eft-3p::Mos1* transposase) at 50 ng/μl and pBluescript at 50 ng/μl. A strain containing the insertion was obtained following standard selection and PCR confirmation ^58^.

*frSi14* and *frSi15* are single copy insertions on chromosome I at the location of the Nemagenetag*Mos1* insertion *ttTi4348* of the pSO9 (*vit-6p::OspF::unc-54 3’UTR*) and the pS010 (*col-19p::OspF::unc-54 3’UTR*) plasmids, respectively. They were obtained by CRISPR using a self-excising cassette (SEC) ^59^ that was modified for single insertions on chromosome I. pSO9 and pSO10 were obtained by insertion of either the intestinal-specific promoter *vit-6* or the epidermis-specific promoter *col-19* fused to the gene for the *Shigella flexneri* protein effector OspF that irreversibly dephosphorylates p38 ^25^ into the pSO5.3 vector. pSO5.3 was made from a vector containing the SEC for single insertion on Chromosome II at the position of *ttTi5605* (pAP087, kindly provided by Ari Pani), changing both homology arms to target chromosome I at the position of *ttTi4348*. pSO9 and pSO10 were injected in N2 at 10 ng/μl together with pDD122 (*eft-3p::Cas9*) at 40 ng/μl, pCFJ90 (*myo-2p::mCherry*) at 2.5 ng/μl, pCFJ104 (*myo-3p::mCherry*) at 5 ng/μl, and pCZGY2747 (sgRNA ttTi4348) at 30 ng/μl (kindly provided by Yishi Jin). Non-fluorescent roller worms were selected then heat shocked to remove the SEC by FloxP as described in ^59^. Plasmid and primers sequences are available upon request.

### Infection

Eggs prepared by the standard bleach method were allowed to hatch in 50 mM NaCl in the absence of food at 25 °C. Synchronized L1 larvae were transferred to NGM agar plates spread with *E. coli* OP50. N2 worms were cultivated at 25 °C until the young adult stage (45 hours) and then transferred to plates spread with *S. marcescens* Db10 which had been cultured in liquid LB at 37 °C overnight, put on NGM plates for 24 hours at 25 °C, then cultured another 20 hours at 25 °C as described ^4^. For infection of *D. coniospora*, L1 larvae were cultivated at 25 °C until the L4 stage (40 hours) before being exposed to fungal spores on standard NGM plate spread with OP50, as previously described ^60^. For infection upon starvation, worms were harvested at the young adult stage, starved for 14 hours on NGM plates with carbenicillin then infected for 6 hours. To observe the effect of infection on the translocation of DAF-16, TJ356 worms were first infected for 13 hours then heat-shocked for 30 min at 37 °C and observed directly and then again after a period of one hour at 25 °C. To count the number of spores, young adult worms were infected for 8 hours at 25 °C then observed on a slide under the microscope. Statistical significance was determined using a two-tailed unpaired student’s T-test (Graphpad Prism).

### qRT-PCR

RNA samples were obtained following Trizol (Invitrogen)/ chloroform extraction and then diluted in nuclease-free water to a final concentration of 100 ng/μl. One μg of total RNA from infected or non-infected worms were used for reverse transcription (Applied Biosystems^™^). Quantitative real-time PCR were performed using 1 μl of cDNA in 10 μl of SYBR Green Applied Biosystem^™^ and 0.1 μM of primers on a 7500 Fast Real-Time PCR System using *act-1* as a control. All primers were designed to be specific for the cDNA (see Table S3 for the sequences). Normalized threshold cycle values were used to calculate fold increases or decreases of RNA levels in samples from test condition compared with controls. The *Ct* values were normalized to changes in *act-1* that were found to not change with infection. The details of the analysis procedure are described in ^61^. Mean and SEMs were calculated from a minimum of 3 independent experiments, statistical significance was determined using a one-tailed ratio paired student’s T-test (Graphpad Prism).

### Fluorescent reporter analyses

Analysis of *nlp-29p::GFP* or *ins-11p::GFP* expression was quantified with the COPAS Biosort (Union Biometrica; Holliston, MA) as described in ^29^. In each case, the results are representative of at least 3 independent experiments where more than 70 worms were analyzed. The fluorescent ratio between GFP and size is represented in arbitrary units. Statistical significance was determined using a non-parametric analysis of variance with a Dunn’s test (Graphpad Prism). Fluorescent images were taken of transgenic worms mounted on a 2 % agarose pad on a glass slide anesthetized with 5 mM levamisole, using the Zeiss AxioCam HR digital colour camera and AxioVision Rel. 4.6 software (Carl Zeiss AG).

### Egg laying and survival analysis

20 to 30 young adult (L4 + 12 hours) IG1564 (*wt; frSi2[col-19p::GPA-12\**]) and IG1647 *(ins-11(tm1053); frSi2[col-19p::GPA-12*])* worms were transferred to fresh NGM plates every 2 hours for 3 times over the course of the first day, with a special care being taken to put them inside the OP50 lawn. F1 worms were counted and analysed for their size using the COPAS worm sorter after 3 days at 25 °C. 15 independent plates were analyzed for each strains. The number of F1 per hour and per adult was calculated for each plate. For survival analysis, the same young adult worms were used and transferred every day until they finished to lay eggs (no FUDR was used), then observed each day until they died ^60^. Statistical significance was determined using a non parametric two tailed student’s T-test for the number of F1 progeny and their length and a Mantel-Cox log-rank test for survival (Graphpad Prism).

### RNA-seq analysis

Total RNA from synchronized young adult worms (N2 and *ins-11(tm1053)* worms, with or without the *frSi2* transgene) was obtained following Trizol (Invitrogen)/ chloroform extraction. The RNA sample was treated with DNAse using RNA clean up kit (Qiagen) then sent for RNA-sequencing (Illumina SE50, 20M reads/sample at Novogene, Beijing, China). The data analysis was done through an in-house standard analysis workflow for RNA-seq. In brief, clean reads were obtained by removing containing adapter sequences, reads containing poly-N and low quality reads. All the downstream analysis was done based on the clean reads with high quality. The single-end clean reads were then aligned to the reference *C. elegans* genome (genome sequence WS235; genome annotation WS260). The reads number mapped to each gene were counted and the expected number of fragments per kilo base of transcript sequence per millions base pairs sequenced of each gene was calculated based on the length of the gene and read count mapped to this gene. Finally, differential expression analysis was performed to determine the differentially expressed genes based on the selected P-value. Differentially-expressed genes were defined using the previously described algorithms ^4^. Genes were annotated using SimpleMine WS260, analysed using Venny ^62^, and class enrichment analyses used YAAT http://bioinformatics.lif.univ-mrs.fr/YAAT/ (Thakur et al. in preparation).

## Acknowledgements

We thank Annie Boned, Gwenaelle Bouget, Martin Fan, Emily Mills and Maxime Bulle for their contributions, Laurence Arbibe for the OspF construct, Ari Pani for the original SEC single insertion vector (pAP087) and CRISPR guide on Chr II, Yishi Jin for CRISPR guide on Chr I, Anne Brunet, Orane Visvikis and members of the lab for discussion. Worm sorting was performed using the facilities of the French National Functional Genomics platform, supported by the GIS IBiSA and Labex INFORM. This work was supported by institutional grants from AMU, INSERM and CNRS and ANR program grants (ANR-12-BSV3-0001-01, ANR-16-CE15-0001-01, ANR-11-LABX-0054 (Labex INFORM) and ANR-11-IDEX-0001-02 (A*MIDEX)). Some nematode strains were provided by the CGC, which is funded by NIH Office of Research Infrastructure Programs (P40 OD010440), or by the National Bioresource Project coordinated by S. Mitani. All authors participated in conducting experiments and analyzing and interpreting data. NP and JE wrote the manuscript. All authors approved the final version of the manuscript.

---

**Figure S1.**
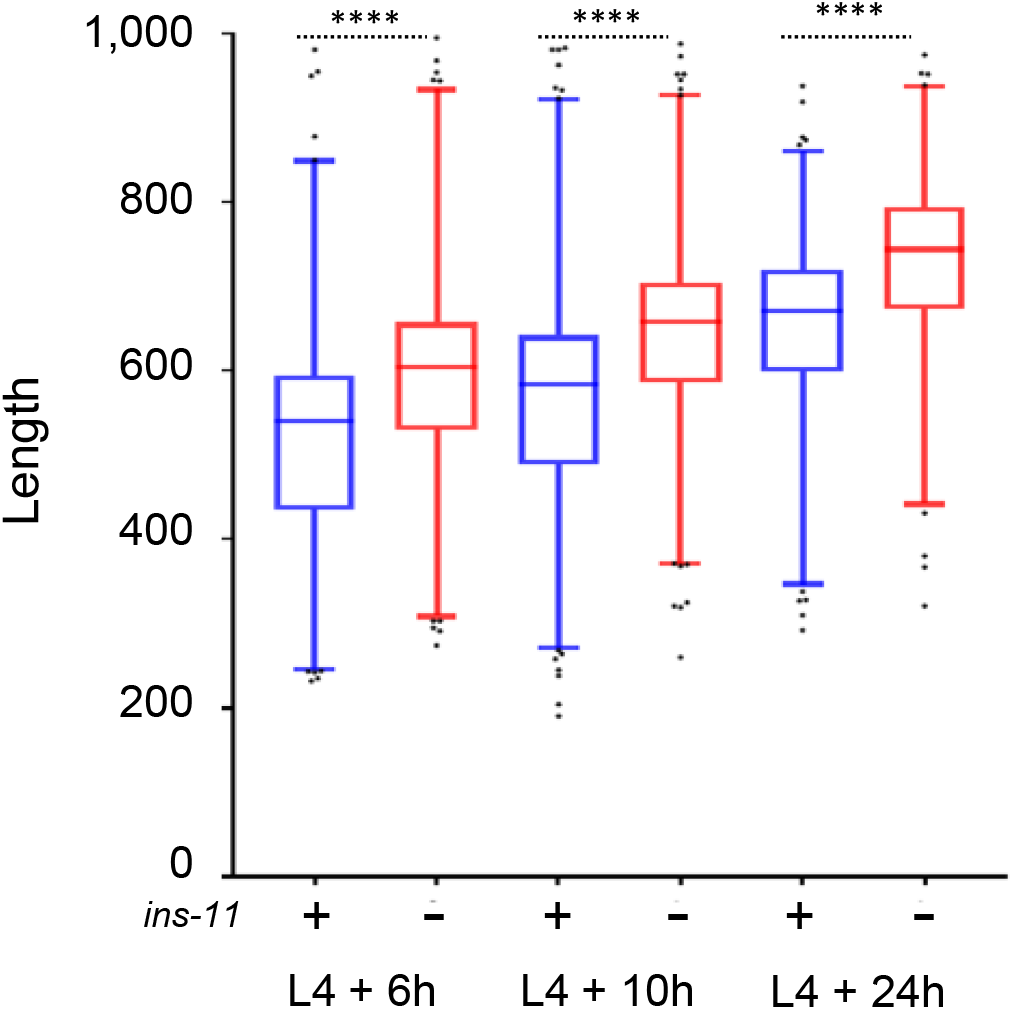
Relative size of young adult worms in wild type (blue) or *ins-11* mutants (red) 6, 10 or 24 hours after the L4 stage (n = 511, 552, 746, 713, 540, 470). Box and whisker plots with 1-99 percentiles, p value < 0.0001.

